# Redesigning chromosomes for optimized Hi-C assay provides insights on loop formation and homologs pairing during meiosis

**DOI:** 10.1101/169847

**Authors:** Muller Héloïse, Scolari F. Vittore, Mercy Guillaume, Agier Nicolas, Aurèle Piazza, Lazar-Stefanita Luciana, Descorps-Declere Stephane, Espeli Olivier, Llorente Bertrand, Fischer Gilles, Mozziconacci Julien, Koszul Romain

## Abstract

In all chromosome conformation capture based experiments the accuracy with which contacts are detected varies considerably because of the uneven distribution of restriction sites along genomes. In addition, repeated sequences as well as homologous, large identical regions remain invisible to the assay because of the ambiguities they introduce during the alignment of the sequencing reads along the genome. As a result, the investigation of homologs during meiosis prophase through 3C studies has been limited. Here, we redesigned and reassembled in yeast a 145kb region with regularly spaced restriction sites for various enzymes. Thanks to this Syn-3C design, we enhanced the signal to noise ratio and improved the visibility of the entire region. We also improved our understanding of Hi-C data and definition of resolution. The redesigned sequence is now distinguishable from its native homologous counterpart in an isogenic diploid strain. As a proof of principle, we track the establishment of homolog pairing during meiotic prophase in a synchronized population. This provides new insights on the individualization and pairing of homologs, as well as on their internal restructuration into arrays of loops during meiosis prophase. Overall, we show the interest of redesigned genomic regions to explore complex biological questions otherwise difficult to address.

## Introduction

Genomic derivatives of the capture of chromosome conformation assay (3C, Hi-C, Capture-C)(Lieberman-Aiden *et al*, 2009; Dekker *et al*, 2002; Hughes *et al*, 2014) are widely applied to decipher the average intra- and inter-chromosomal organization of eukaryotes and prokaryotes (Sexton *et al*, 2012; Le *et al*, 2013; Dekker *et al*, 2013; Marbouty *et al*, 2014a). Formaldehyde cross-linking followed by segmentation of the genome by a restriction enzyme (RE) are the first steps of the experimental protocol. The basic unit of “C” experiments therefore consists of restriction fragments (RFs) that are subsequently religated and captured to identify long range contacts. The best resolution that can be obtained is directly imposed by the positions of the RE sites along the genome. Both 6-cutter and 4-cutter REs have been used (Marie-Nelly *et al*, 2014; Sexton *et al*, 2012; Rao *et al*, 2014; Le *et al*, 2013), the latter with the expectation that the resolution increases with the number of sites. However, this approach suffers from two major caveats. First, restriction sites (RSs) are not regularly spaced along genomes. The distribution of RFs lengths follows a geometric distribution, with important variations along the genome that depend on the local GC content and the specific sequence recognized by the RE. Given that the likelihood for a RF to be crosslinked by formaldehyde during the first step in the procedure depends on its length (Cournac *et al*, 2012), the probability to detect a given fragment in any 3C experiment will in turn be strongly affected by this parameter (**Fig 1A**). Normalization procedures have been developed in order to correct the signal (Cournac *et al*, 2012; Imakaev *et al*, 2012) but these methods involve filtering out fragments with unusually low or high signal and aggregating the contact data over several consecutive fragments in longer bins of fixed genomic length, at the expense of actual resolution (Lajoie *et al*, 2015). Overall, the definition of Hi-C resolution has remained empiric, because of the lack of a control sequence where RF biases would be alleviated. The second limitation reflects the fact that repetitive sequences cannot be tracked because the sequencing reads obtained from a Hi-C experiment cannot be mapped unambiguously along the genome, alleviating the possibility to track homologous chromosomes in isogenic backgrounds.

**Figure 1.**
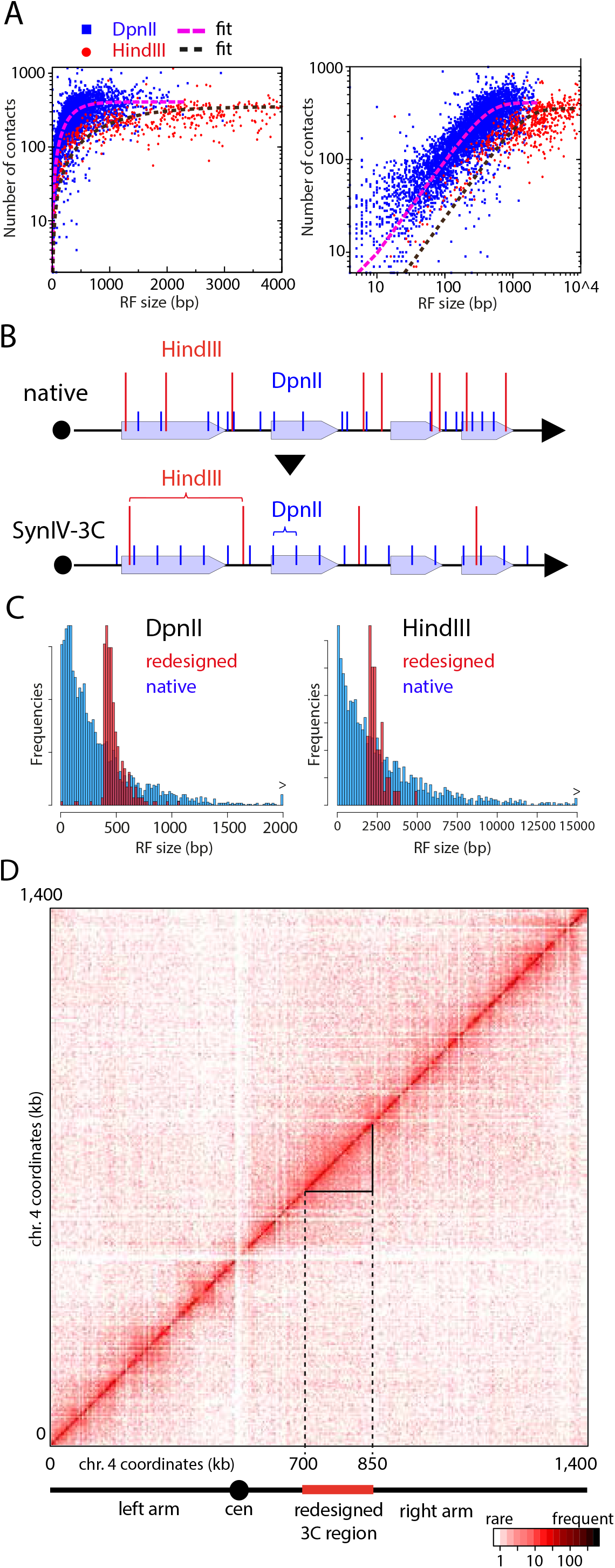
synIV-3C design and assembly. A Number of contacts made by RFs as a function of their size (HindIII (red) or DpnII (blue) in the native sequence. Left panel: log-lin scale. Right panel: log-log scale.). B Illustration of the design principles of the synIV-3C sequence for the DpnII and HindIII RSs. Black arrow: chromosome. Grey rectangles: genetic elements. Blue and red vertical lines represent the RSs positions for the enzymes DpnII and HindIII, respectively. Top panel: restriction pattern of a (hypothetical) native sequence. Bottom panel: restriction pattern after synIV-3C design, with the RSs defining regularly spaced intervals. C Distribution of the DpnII (left) and HindIII (right) RFs sizes in both the native and synIV-3C 150kb redesigned sequence (red and blue, respectively). D Raw DpnII contact map of the Hi-C experiment performed on G1 daughter cells synchronized through elutriation (Marbouty *et al*, 2014). Dashed lines: borders of the redesigned region. Plain black lines: borders of the contact map analyzed in **Fig. 2**.

**Figure 2.**
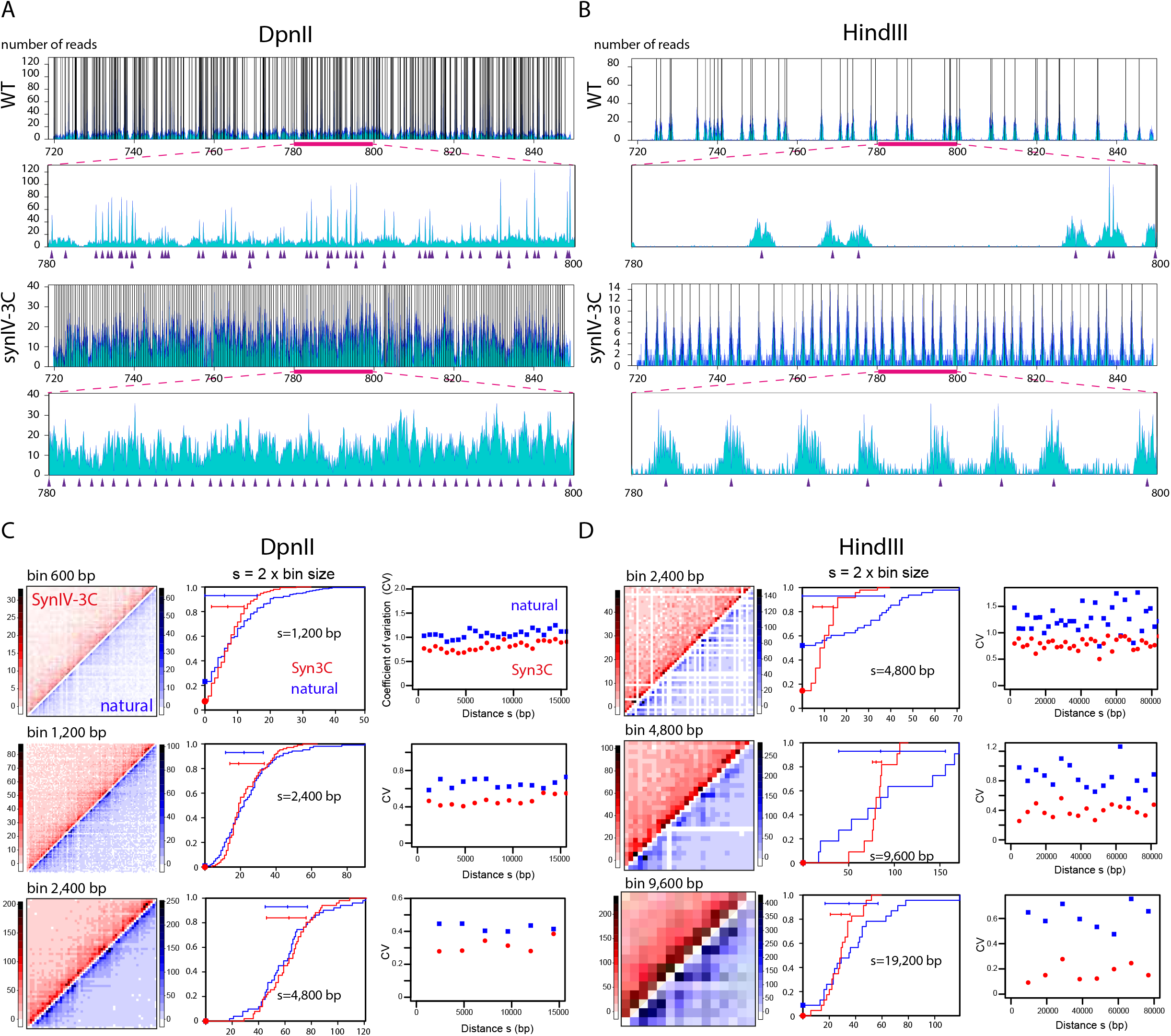
Sequence reads coverage from Hi-C experiments performed with DpnII (A) and HindIII (B) restriction enzymes in synIV-3C and native strains, and mapped against the synthetic region SynIV-3C and its natural WT counterpart, respectively. For each region, the magnification of a 20 kb window (pink line) is shown. Vertical lines indicates the position of restriction sites in the top panel and purple triangles their position in the magnified 20kb region. Note that the scale of the y-axis illustrates the heterogeneity of the coverage, with some positions in the DpnII map being overrepresented with respect to others. C, D Analysis of the contact counts along the synIV-3C region for DpnII (C) and HindIII (D). First column: synIV-3C (in red) and chromosome 4 native counterpart (blue) Hi-C contact maps. For each experiment, three different fixed bin sizes were analyzed (600 bp, 1200 bp and 2400 bp for DpnII, 2400 bp, 4800 bp and 9600 bp for HindIII). Middle panels: cumulative distribution of the number of contacts between bins located at a genomic distance ***s*** from each other’s (***s*** = 2 x bin size). Right panels: distribution of the coefficient of variation (CV) as a function of ***s***.

One consequence of these limitations has been the absence of in-depth studies of meiotic prophase through Hi-C approach. Meiosis is the cell division where a diploid cell gives rise to four haploid gametes through two rounds of chromosome segregation with no replication in-between them. The prophase of the first division, where the homologous paternal and maternal chromosomes segregate, comprises a series of regulated events involving the recognition and pairing of homologs all along their length. Homologs, to become physically connected, must meet each other’s in space, which implies a dynamic reorganization of the overall genome and disentanglement between paired DNA molecules (Zickler & Kleckner, 2016). During the meiotic program shared by budding yeast and mammals (i.e. the succession of events mediated by evolutionary conserved molecular complexes), pairing occurs early on during the leptotene stage. This process can be accompanied and/or facilitated by dynamic movements of chromosomes, and lead to telomere clustering at the zygotene stage (bouquet stage; (Zickler & Kleckner, 2016) or other forms of movements mediated by chromosome ends directed by cytoskeletal components through direct association across the nuclear envelop (Koszul & Kleckner, 2009). These events have mostly been described using imaging techniques, or, when it comes to the analysis of the underlying molecular events, through site-specific assays. Dekker and coworkers pioneered the analysis of meiosis in the original chromosome conformation capture study (Dekker *et al*, 2002). Using restriction polymorphism to distinguish the maternal and paternal versions of a locus along chromosome III, they notably showed that 3C was able to capture homolog pairing, as well as centromere declustering. However, the higher-order organization surrounding the recombining locus remained unexplored. Similarly, the influence on homolog pairing of the vigorous movements mediated by chromosome ends remain unexplored using Hi-C (Koszul & Kleckner, 2009).

In order to investigate the behavior of two homologous chromosomes sharing the same nuclear space, we designed and assembled a dedicated “synthetic” genomic region (Koszul, 2016), also aimed at increasing the resolution of 3C-based experiments. As a proof of concept of this strategy, we describe here a redesigned ~150kb region (called synIV-3C) of *Saccharomyces cerevisiae* yeast chromosome 4. We then investigated its behavior during the first stages of meiotic prophase.

## Results and Discussion

### Design and assembly of the Syn-3C region

Designer chromosome synIV-3C closely resembles the native chromosome with respect to genetic elements (see Material and Methods), but was “designed” to yield high resolution and high visibility in 3C experiments by providing nearly equally spaced restriction sites. The RSs of four different enzymes were removed from the native sequence with point mutations and subsequently reintroduced within the sequence at regularly spaced positions (400bp, 1,500bp, 2,000bp and 6,000bp for DpnII, XbaI, HindIII and NdeI, respectively; **Fig 1B** and **Fig S2**; **Table 1**). As shown on **Fig 1C**, the DpnII and HindIII RFs sizes in the redesigned synIV-3C region are normally distributed when compared to the skewed, native genome-wide distributions. Besides providing a way to increase the resolution of the 3C experiment, the design can also be used to focus on specific contacts, for instance between promoters and terminators (**Fig S2**). When possible, coding sequences were targeted preferentially and modified using synonymous mutations (**Fig S1**). We identified a 150kb window on chromosome 4 for which the uniformity of RFs lengths was maximized while the number of potentially deleterious base changes was minimized (the final choice for the region can also take into account sequence annotation and be guided by specific interests of the end-user). From this design, DNA building blocks were purchased and assembled in yeast BY (S288C) and SK1 background strains as described (Annaluru *et al*, 2014; Muller *et al*, 2012) (Material and Methods). Sequencing confirmed that 144kb within the targeted region were replaced by the redesigned sequence and that 100% of the mutations were introduced at the correct positions corresponding to a total of ~2% divergence with the reference genomes (3,229 bp out of 150,000). Analysis of the growth profile did not reveal significant negative effects of the modifications introduced in the SynIV-3C region compared to the isogenic parental strains (**Fig S3**).

**Table 1.** Mutations necessary to remove and generate new sites along chromosome 4 700,000::850,000 window.

|  | deletion | new sites |
| --- | --- | --- |
| HindIII | 58 | 61 |
| NdeI | 34 | 23 |
| XbaI | 25 | 76 |
| DpnII | 442 | 310 |
| **Total** | 559 | 470 |

### *Cis*-contact pattern of the SynIV-3C region

To assess for the quality improvement of Hi-C data in the Syn-3C region, Hi-C experiments were performed in parallel on BY strain carrying the synIV-3C redesigned chromosome as well as on the native parental strain using DpnII and HindIII (Material and Methods). The raw DpnII contact map of chromosome 4 exhibited a remarkably “smooth” pattern within the redesigned region compared to the native flanking regions (**Fig 1D**). The read coverage over the region also exhibits a dramatic and compelling change, with a more homogeneous and regular distribution in the synthetic regions for both enzymes compared to a heterogeneous distribution in the native sequence (**Fig. 2A, B**). Interestingly, careful examination of this distribution indicates that besides its own length, the capture frequency of a given fragment is also influenced by the length of its neighbors, resulting in a bias. To quantify the improvement in the SynIV-3C region we compared the signal with the signal over the same region obtained in the WT strain using the same number of aligned read pairs and identical bins of various sizes (**Fig. 2C, D**). At the smallest resolution tested (600bp for DpnII and 2,400bp for HindIII) the WT contact map exhibited numerous blind regions with no detectable contacts (empty bins), in sharp contrast with its synthetic counterpart (**Fig. 2C, D**). When fragments were aggregated in bins of increasing sizes (hence, resulting in a loss of resolution) these blind regions gradually disappear, although the heterogeneity of the data remains consistently higher in the WT compare to synIV-3C strain, as shown by the increased span of the color-scales of the WT maps.

In order to further quantify this heterogeneity, we computed the cumulative distributions of the number of contacts between bins separated by a given genomic distance ***s*** (bp) in the synIV-3C region and in its native counterpart for DpnII and HindIII (**Figs 2C** and **2D**, respectively). The redesigned region systematically exhibited more homogeneous contacts counts and narrower distributions than the WT region, both at short (*s* = 2 x bins sizes; **Figs 2C** and **D** middle panels) and longer distances (Material and Methods and **Fig. S5, S6**). Some of the bins in the native region remain almost invisible to the assay as a result of the heterogeneity in RF distribution (blue squares on **Figs 2C** and **D** middle panels). We computed the coefficient of variation CV (i.e. standard deviation /mean) of these distributions for multiple values of *s*. We use this value as an indication of the signal to noise ratio (**Figs 2C** and **D** right panels). Interestingly, we found that even for large bins, the CV is significantly and consistently smaller in the synthetic region, again indicating improved resolution. These results also clearly illustrate the advantage of using a frequent cutter (DpnII vs. HindIII) restriction enzyme with respect to resolution since the distribution of contact counts between bins remains much more spread with HindIII than with DpnII, even for native sequences (**Fig 2B**).

### Statistical analysis of Hi-C contact data

The sequencing step of a Hi-C library corresponds to the random sampling of all ligation events generated during the experiment. The outcome of this draw, for a given genomic distance s ± Δs between pairs of loci, is expected to follow a Poisson distribution if we suppose that each pair of loci has a strictly equal probability p to be drawn. For real data, this cannot be the case since other factors (such as for instance the restriction fragment size, or the average 3D folding of chromatin) can influence this probability. When the number of events becomes large enough though, the differences between the re-ligation probabilities of different loci will start to kick in and the distribution of contacts should switch to a Gaussian behavior, assuming that these probabilities follow a Gaussian distribution. The standard deviations (σ) of those two distributions (Poisson and Gaussian) scale either as the square root of the mean (μ) or as the mean, respectively. We took advantage of the fact that the overall contact number μ decreases with increasing genomic distances (s) to check whether these relations between μ and σ holds for real data (**Fig. 3A**).

**Figure 3.**
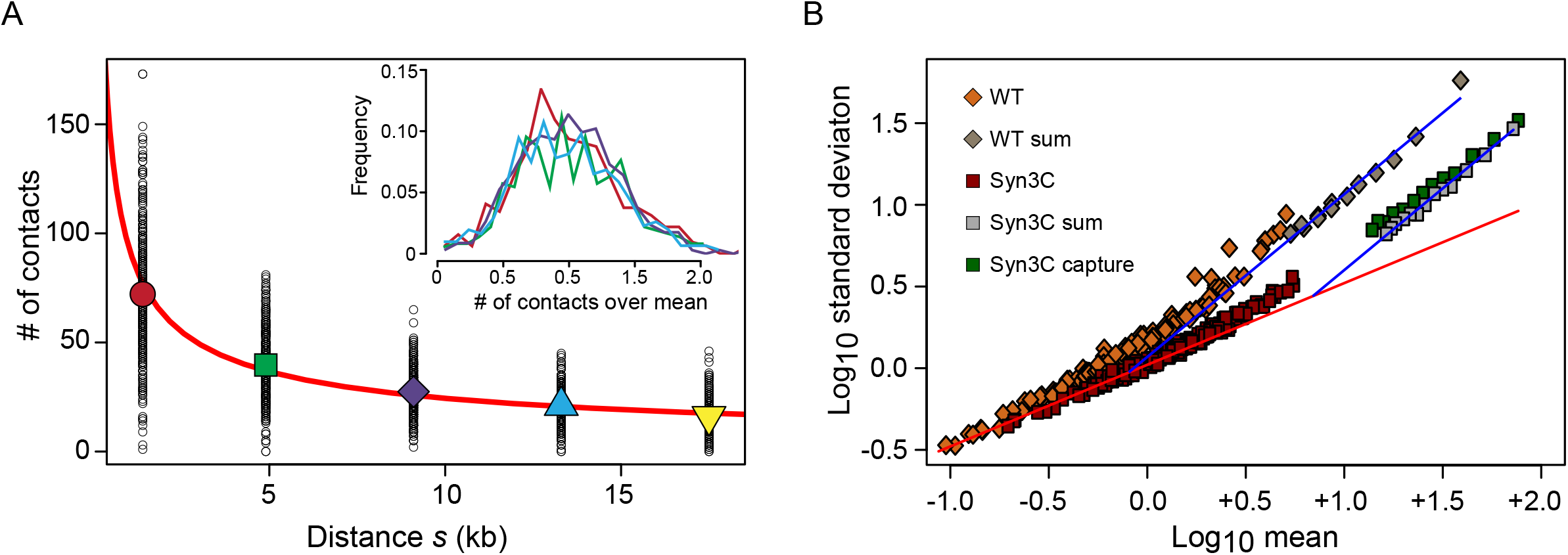
Relation between standard deviation and mean for each pair of fragments located at similar genomic distances s ± Δs. A The mean number of contact of restriction fragments (big symbols) at a fixed range of distances (s +/- 350bp, see text) decreases as a power law (red line) with exponent -0.6. Small circles represent the contact number of individual pairs of restriction fragments. Inset: Distributions of contact numbers re-scaled by their means B The relation between the variance and the mean of these distributions undergoes a transition from Poisson to Gaussian. WT and Syn3C data were obtained by aggregating data-sets respectively from (Mercy *et al*, 2017) and (Lazar-Stefanita *et al*, 2017). Capture-C data are from this study. The red line corresponds to the theoretical Poisson behavior, whereas the blue line corresponds to the theoretical Gaussian behavior fitting the CV from data.

To start our analysis with the highest contact numbers we aggregated the results from 12 Hi-C experiments performed in G1 or early S phase (from Lazar-Stefanita *et al.*, 2017). In the following, we focused our analysis on the Syn3C region. For five different distances (**Fig. 3A**; 1,400 ± 350 bp in red; 4,900 ± 350 bp in green; 9,100 ± 350 bp in purple; 1,330 ± 350 bp in cyan and 17,500 ± 350 bp in yellow), the five contact distributions computed from each restriction fragments pairs were re-scaled by their mean and superposed on the same plot (**Fig. 3A**, inset). The collapse of these re-scaled distributions clearly indicates a Gaussian behavior (i.e., the mean scales as the standard deviation). We next explored the relation between μ and s over a wider range of values. We used data for increasing values of s, corresponding to lower μ, as well as data aggregated over multiple experiments and data from capture experiments, corresponding to higher values of μ (**Fig. 3B**). When plotting the values of s for different μ, we found that both in the native and in the Syn-3C sequence context there is a crossover from the Poisson to the Gaussian distribution (indicated by the red and blue lines, respectively) and that the standard deviation for the Syn3C experiment in lower than in the WT counterpart for all the values of μ, as expected from the analysis done in **Fig. 2**. Interestingly, the two transition points can tell us about the importance of the bias of having uneven restriction fragments compared to other biases and/or biologically relevant variations. In the WT case, as soon as each pair of fragments receive one read, the distribution of counts switches to Gaussian, indicating that each pair has already been sampled unevenly. In the case of the Syn3C construct, where this bias is absent, one needs to aggregates 10 reads per fragment pairs to start to see variations among fragment pair re-ligation frequencies and switch to the Gaussian behavior. This highlights the strong effect of fragments length biases and justifies the use of large bins which will encompass many fragments as well as normalization procedures in all Hi-C experiments. For any genomic distance s, the value of μ can arbitrarily be increased by increasing the bin size, or resolution.

The existence of the transition between Poisson and Gaussian behaviors thus enables us to propose a rigorous way to determine the resolution of a Hi-C experiment by choosing a bin size which exactly corresponds to the transition point. It is worth underlying that according to this analysis the resolution of a Hi-C experiment can only be defined for a given genomic distance s.

### Analysis of genome organization during meiosis prophase

The SynIV-3C chromosome was designed and assembled with the aim to investigate the interplay between homologous chromosomes during a variety of DNA related metabolic processes, including the mitotic and meiotic cell cycle. A diploid SK1 strain carrying the SynIV-3C region on one homolog, its native counterpart on the other homolog, but isogenic for the rest of the genome, was processed into a synchronized meiotic culture (Material and Methods). The synchrony of meiotic progression was assessed by monitoring meiotic replication by FACS analysis and the two meiotic divisions by DAPI staining. Cells that have passed through anaphase I or anaphase II contain two or four DAPI-stained bodies, respectively. After 6h in sporulation medium, ~40% of the cells display two or more DAPI bodies. By 8h, 70% of the cells are passed anaphase II, showing that most cells have completed meiosis synchronously (**Fig. 4B**) (Hunter & Kleckner, 2001; Koszul *et al*, 2008). Hi-C contact maps were generated for cells sampled at 0, 3 and 4h from the synchronized culture, corresponding to WT mitotic, early zygotene and early pachytene cells (when a fraction of the population has passed anaphase I), respectively (**Fig. 4C**). The differences between mitotic and early pachytene (4h) cells were determined by computing the log-ratio between the contact maps (bin: 5 kb; **Fig. 4D**; Material and Methods). Under this representation, the color scale of the map reflects the variations in contact frequency for each bin between two different contact maps. Individual 2D maps can also be represented as 3D structures to facilitate their interpretation (**Fig. 4E**; Lesne *et al*, 2014; Lazar-Stefanita *et al*, 2017). These different representations illustrate and recapitulate known key features of meiotic prophase.

**Figure 4.**
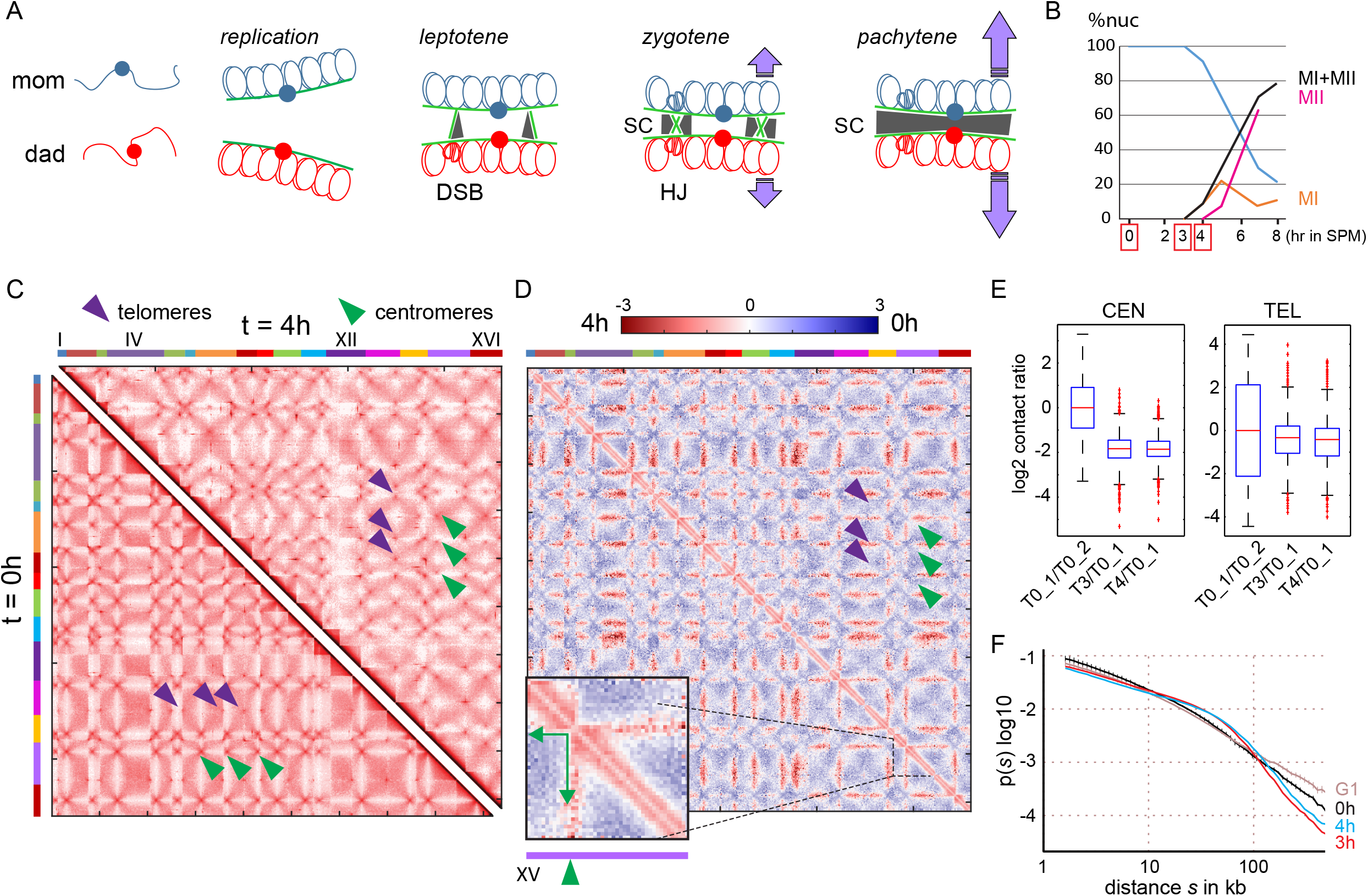
Individualization of chromosomes during prophase. A Schematic representation of the structural changes affecting homologs during meiotic prophase. Sister chromatids organize as arrays of loops along each homolog axis (green lines) after replication. At leptotene, DNA double-strand breaks (DSBs) occur. During zygotene, homologs come together partially and the syneptonemal complex (SC, grey) originates at DSBs and centromeres. At pachytene, homologs are fully synapsed and undergo vigorous motion (purple arrows). B Meiotic progression, as measured by completion of MI and MII divisions. C Contacts maps of synchronized populations of cells after 0 (bottom left) and 4 (top right) hours in sporulation medium. The 16 yeast chromosomes are displayed atop the maps. Purple arrowheads: inter-telomere contacts. Green arrowheads: inter-centromeric contacts. D Log-ratio of the two contact maps from (C). The blue to red color scale reflects the enrichment in contacts in one population with respect to the other (log2). The inset on the bottom left displays the magnification of chromosome XV. The green arrows point at the contacts made at t=4h by the centromere with the left and right arms. E Boxplots representing the variation in number of normalized contacts for centromeres (left) and telomeres (right) between 0, 3 and 4h in sporulation media. Variations between t=0h replicates, 3h and 0h, and 4h and 0h, were computed. The relative Wilcoxon test supports a decrease in centromere contacts at t=3 and 4h compared to t=0h. F Average intra-chromosomal contact frequency p between two loci with respect to their genomic distance s along the chromosome (log-log scale; p(s)) during mitotic G1 (brown curve; three replicates from Lazar-Stefanita et al., 2017), meiotic t=0h (black curve; two replicates), 3h (red curve) and 4h (blue curve).

#### Declustering of centromeres

A loss of inter-centromeric contacts was readily apparent during zygotene and pachytene compared to the pre-replication stage (green arrowheads; **Fig. 4C, D, F**). No significant enrichment in contacts between telomeres was observed (purple arrowheads; **Fig. 4C, D, F**). This result reflects the rapid declustering of centromeres that accompanies entry into meiotic prophase also observed through microscopy and 3C (Dekker *et al*, 2002; Trelles-Sticken *et al*, 1999). On the other hand, the absence of inter-telomeric contacts doesn’t immediately support the transient and small increase in telomere clustering described at this stage and corresponding to the bouquet stage ((Zickler & Kleckner, 1999; Trelles-Sticken *et al*, 1999). One possibility is that the subtelomeric sequences of the SK1 strain remain incomplete or incorrectly assembled and therefore escape monitoring using Hi-C. Alternatively, the transient increase in clustering is very rapid, or occurs a bit earlier, and was missed during this experiment. More experiments are therefore necessary to fully assess whether or not budding yeast meiotic program really present a “bouquet” stage.

In addition to the loss of discrete centromeric contacts, the ratio map also revealed the abolition of the “polymer brush” effect, which insulates centromeres and their flanking chromosomal regions from the rest of the chromosome arms (green arrows on the magnification of chromosome XV, **Fig. 4D**). As a result, the centromeric regions become almost indistinguishable from the rest of chromosome arms, and chromosomes appear as relatively individualized, homogenous entities. Once declustered, centromeres regions are more likely to contact other portions of the genome, resulting in the increased *trans* contact signal made by these regions with other chromosomes and visible on the ratio map (**Fig. 4D**). The transformation of chromosomes into well-individualized entities could be strikingly recapitulated by the 3D representation of the 2D maps (**Fig. 4E**) (Lesne *et al*, 2014).

#### Chromosome folding

The ratio contact maps also display a strong increase in intra-chromosomal contacts at early-pachytene compared to mitotic cells, as reflected by a red and large diagonal that clearly appears when magnifying individual chromosomes (inset on **Fig. 4D**). The folding can also be assessed by computing the contact probability *p* as a function of genomic distance of all chromosome arms (Naumova *et al*, 2013; Lieberman-Aiden *et al*, 2009; Lazar-Stefanita *et al*, 2017). The p(s) curves were computed for two premeiotic replicates (0h), three mitotic G1 replicates (Lazar-Stefanita et al., 2017), zygotene (3h) and early-pachytene (4h) chromosomes. The two later curves display sharp differences compared to premeiotic and G1 mitotic cells. First, contacts frequencies increase between loci positioned 20 to 50 kb apart, with a peak around 50 kb (**Fig 4G**). Then, contacts sharply decrease, suggesting a general stiffening of the polymer associated with the loss of distant loci to contact each other’s. This pattern suggests that chromosomes fold during meiotic prophase into a structure that favor contacts under a certain distance, while disfavoring local and long range interactions, and points at the formation of chromatin loops.

### Visualization of meiotic loops using Hi-C

After or during meiotic replication, the two sister chromatids of each homolog becomes organized as arrays of ~20 kb loops tethered to a protein structural axis composed (**Fig. 3A**) (Zickler & Kleckner, 2016; Blat *et al*, 2002; Zickler & Kleckner, 1999). The meiosis-specific cohesion subunit Rec8 is one of the component present along these axis (Klein *et al*, 1999). Rec8 is distributed into discrete domains corresponding to meiotic loops bases, and contributes to the establishment of cohesion between sister chromatids. These loops provide a highly-organized chromosome architecture context for the formation and resolution of meiotic DNA doublestrand breaks generated during leptotene (Padmore *et al*, 1991; Blat *et al*, 2002; Kim *et al*, 2010; Sommermeyer *et al*, 2013; Acquaviva *et al*, 2013). It was proposed that the expansion state of the chromatin within the loops modulates the degree of compaction and stiffness of the chromosomes at the different stages of prophase (Kleckner *et al*, 2004; see also Koszul *et al*, 2008). The impact of loop formation on chromosomes had never been addressed through Hi-C, and we took advantage of this pilot study to have a glimpse on these processes.

First, we investigated whether Rec8-mediated loops were visible on the Hi-C contact maps of zygotene and early-pachytene cells by displaying the Rec8 binding regions along chromosome axis. Both at 3 and 4h, the short range intra-chromosomal contacts displayed a heterogeneous signal, with triangular shapes domains reminiscent of the topological domains observed along the mitotic chromosomes of other species appearing along chromosomes (**Fig. 5A**) (Dixon *et al*, 2012; Sexton *et al*, 2012; Nora *et al*, 2012). The boundaries between these domains correlate with Rec8 enriched regions (Ito *et al*, 2014; Glynn *et al*, 2004), suggesting that this signal may correspond to chromatid loops bridged at their bases to the chromosomal axis.

**Figure 5.**
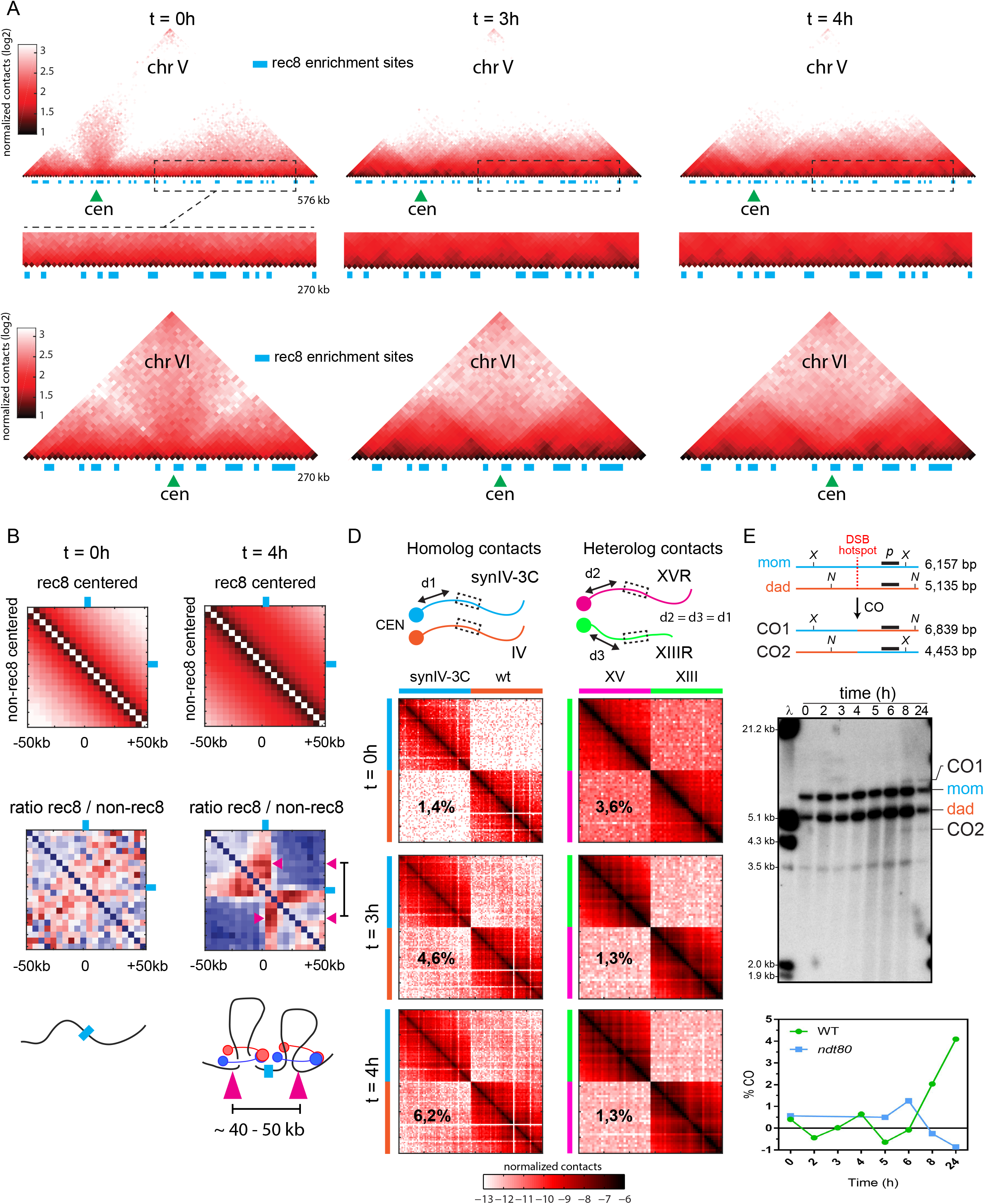
Organization of homologs into chromatin loops. A Magnification of chromosomes V and VI contact maps during meiotic time course at t=0, 3 and 4 hours (bin size: 5kb). The blue rectangles point at bins enriched in Rec8 protein. Green triangles: centromere position. B Top panels: aggregated intrachromosomal contacts made by 100kb windows (5 kb bins) centered on Rec8 enriched bins (top right) or randomly chosen (bottom left). Middle panels: ratio between the cumulated contact maps of the above panel. Blue color shows a depletion of contacts in the random maps, whereas the red signal points at an enrichment in contacts in the maps centered on Rec8 enriched bins. Purple arrowheads point at looping signal between the center of the window (corresponding to the Rec8 enriched bin) and flanking regions. Bottom panel: schematic representation of the disposition of the chromatin at the corresponding timepoints. C Normalized frequency of contacts as a function of genomic distance within the SynIV-3C and its WT counterpart for t=0, 3, and 4h. D Contacts between the SynIV-3C region and its native homolog (left panels) and between two windows of similar sizes positions at an equal distance (d) to the centromere but belonging to two other chromosomes (right panels). The percentages on each column reflects different measurements. For the inter-homolog panels, they represent the number of pairs of reads bridging homologs (*trans* contacts) divided by the total number of pairs aligning within the regions. On the other hand, for inter-heterologs, the percentage represents the amount of pairs of reads bridging the two distinct regions, divided by the total number of pairs aligning within the distinct regions. This illustrates the gradual chromosomal individualization. E Crossing over between the SynIV-3C region and the native counterpart. Top: schematic representation of how the redesigned restriction pattern allows characterization of CO events using a probe (black line p) at a DSB hotspot (Material and Methods). Middle: meiotic recombination of cells progressing into meiosis and analyzed using restriction pattern similar to (Hunter & Kleckner, 2001). Parental homologs, ‘‘Dad’’ and ‘‘Mom’’, and COs are distinguished on Southern blots via restriction site polymorphism. Bottom panel: CO as percent of total hybridizing signal with time after transfer to sporulation medium.

To further test whether these domains are indeed meiotic loops, we compared the aggregated intrachromosomal contacts made by Rec8-binding sites to random positions in mitotic (0 h) and early-pachytene cells (4h). The ratio between the cumulated contact maps made by 100kb windows (5 kb bins) centered on Rec8 binding sites and random maps display no significant enrichment in mitotic conditions (**Fig. 5B**, left column). At 4h into meiotic prophase, the same analysis display a strikingly different pattern, with the Rec8 bound sites now clearly delimiting two distinct domains (**Fig. 5B**, right column). In addition, these sites now display enriched contacts with DNA regions positioned on average 20 to 25 kb upstream and downstream the chromosome. This contact pattern is typical of loops bridging distant loci in *cis* (Rao *et al*, 2014; Lazar-Stefanita *et al*, 2017), and altogether these results demonstrate that Rec8 is positioned at the basis of chromatin loops during meiotic prophase. Therefore, Hi-C detects loop formation during meiotic prophase. The difference between zygotene and early-pachytene stages was not readily apparent from this work, although changes in chromosomes stiffness have been reported and proposed at these stages (Koszul *et al*, 2008; Kleckner *et al*, 2004). One possibility is that at 4h the amount of cells with fully complete synaptonemal complex axis (pachytene cells) remains limited. Another option is that the signal within the loops is so strong that variations over long distances between zygotene and pachytene remain negligible when assessed by Hi-C. More work on synchronous cultures of wild type and mutant cells should answer these questions and provide new insights on these mechanisms.

### Homolog-homolog contacts using the SynIV-3C and native regions

To characterize more specifically the inter-homolog contacts between the synthetic and native chromosome IV regions an enrichment step for this region was performed using a Capture-C strategy (Hughes *et al*, 2014). This led to a ~20 fold increase in reads from the SynIV-3C region and its native counterpart that were used to generate the contact map at 0, 3 and 4 hours (**Fig. 5C** right panels; see Supplementary methods for controls). To monitor whether changes in contacts between the two homolog regions may result from their similar distance to the centromere, the contact pattern of two heterologous 150kb regions positioned at equal distances from the centromeres on arms of approximately similar sizes (XVR and XIIIR) were generated (**Fig. 5C**, left panels). These heterologous regions became more insulated from each other’s as cell enter meiotic prophase, reflecting the individualization of chromosomes and loss of Rabl organization (see also **Fig. 4D**). On the other hand, the SynIV-3C and native homologous regions, which are clearly distinguishable from each other’s thanks to the SNP introduced in the synthetic design, display a contact frequency that increases over time, from 1.4% of reads before replication to 6.2% in early pachytene.

The two homologs do not appear to be closely and/or strongly juxtaposed, as shown by the weak diagonal that appears over time in zygotene and early pachytene cells. This observation strongly suggests that the synaptonemal complex plays a role not only in the maintenance of homolog structure by bridging chromosomal axis, but also in insulated the two homologs from each other’s. To overcome this barrier, meiotic DNA double-strand breaks are indeed relocalized to the SC, where inter-homolog contact is promoted and eventually recombination can proceed (Sommermeyer *et al*, 2013; Acquaviva *et al*, 2013). Because the SynIV-3C and native regions differ on average by 2%, we wondered whether this polymorphism could affect their folding and/or meiotic recombination processes. The computation of the p(s) for each of the two homologous region at the three timepoints reveal a similar trend than the one observed for the whole genome (**Fig. 5D**), showing that the design does not affect chromatin folding. To verify whether this design impairs meiotic recombination, we performed a restriction assay taking advantage of the restriction polymorphism introduced between the two regions (**Fig. 5E**). Crossing over were detected at the meiotic DSB hotspot tested within the region, suggesting that meiotic recombination can proceed in this genetic environment (**Fig. 5E**). More analyses remain to be performed to verify that this is the case over the entire region. Eventually, the synthetic design could be adapted to alleviate this concern, for instance by removing polymorphisms within 2 kb of DSB hotspots of interests. Nevertheless, this experiment shows that the present approach allows to track recombination events concomitantly to the higher-order architecture of the chromosomes.

### Conclusion

The yeast genome presents a relatively homogeneous GC content and a few repeated sequences. The gain in resolution achieved by redesigning RS along the genome should therefore be even higher in organisms with more heterogeneous genomic content and will enable unbiased tracking of entire regions that are otherwise inaccessible to the experiment. One could envision, for instance, assembling the redesigned chromosome in yeast (Benders *et al*, 2010), before replacing its native counterpart in the organisms of interest (such as a bacteria, or eventually on mammalian cells). Other advantages of the approach include the modularity of the assembly step (**Supplementary Note 2**), that allows the introduction of building blocks carrying genetic elements of interest within the redesigned region. For instance, one could introduce highly expressed promoters in the middle of “gene desert” areas, to investigate the effect of gene expression on the local chromatin structure. One can also “shuffle” some of these building blocks, to look at the influence of specific DNA binding proteins on the contact networks. This specific 3C-friendly design is the first time, to our knowledge, where a large (>100kb) region of chromosome is specifically redesigned and assembled for the purpose of improving an assay so that we can now address more precisely and accurately specific questions related to the biology of the cell. It paves the way to more studies exploiting the power of synthetic biology to boost, refine, and maybe reshape traditional molecular biology approaches through orthogonal ones.

Here, we take advantage of the design to track for the first time using Hi-C the large structural changes that affect chromosomes during meiosis prophase. We showed that Hi-C clearly points at the formation of loops, whose bases overlap with binding sites of the meiotic-specific cohesion subunit Rec8. The redesigned region now also allows to distinguish, to a large extent, the two homologs. Preliminary results suggest that the loops along each homolog are not closely juxtaposed in space, and that the synaptonemal complex constitutes a barrier in-between. This observation further supports the mechanical necessity to generate bridges between homologs for recombination to proceed.

## Material and Methods

### Strains

The SynIV-3C region was assembled in the s288C and SK1 genetic backgrounds by transformation of BY4742 and ORT4602 (Sollier *et al*, 2004) to generate strains RSG_Y181 and RSG_Y189, respectively (see above). The SK1 diploid heterozygous for the SynIV-3C region (RSG_Y190) was obtained by crossing RSG_Y189 and ORT4601 (Sollier *et al*, 2004). Genotypes are provided in **Table 2**.

**Table 2.**
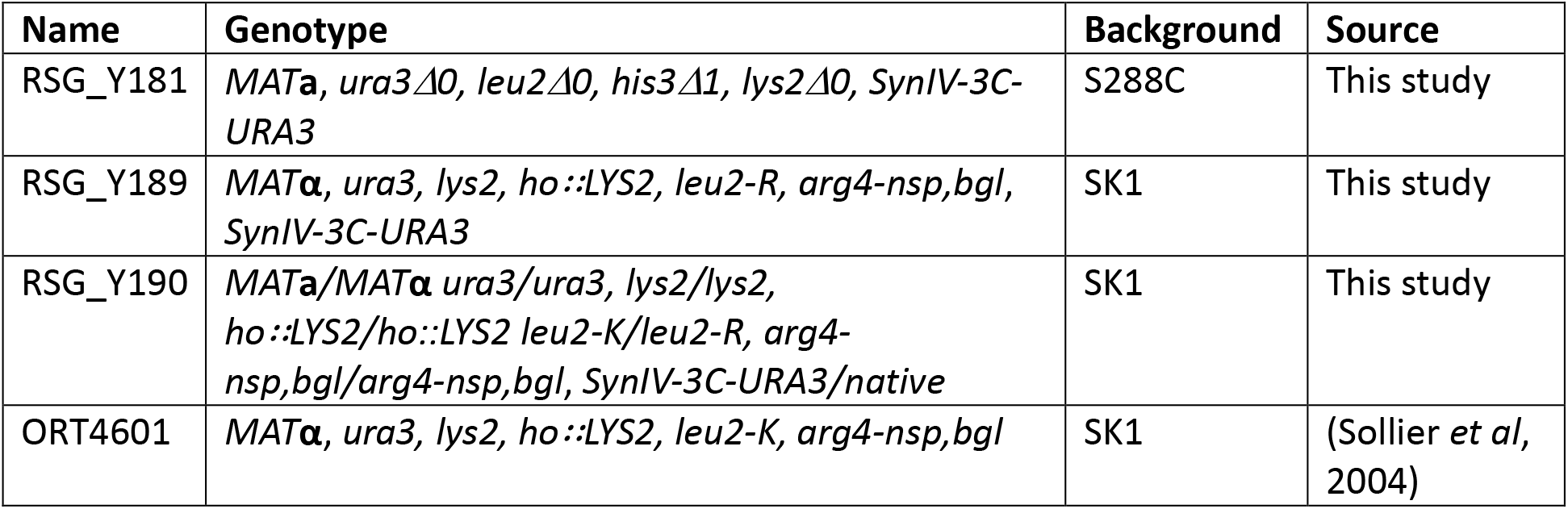
Genotype of *Saccharomyces cerevisiae* strains used in this study.

| Name | Genotype | Background | Source |
| --- | --- | --- | --- |
| RSG_Y181 | *MATa, ura3Δ0, leu2Δ0, his3Δ1, lys2Δ0, SynIV-3C-URA3* | S288C | This study |
| RSG_Y189 | *MATα, ura3, lys2, ho::LYS2, leu2-R, arg4-nsp,bgl, SynIV-3C-URA3* | SK1 | This study |
| RSG_Y190 | *MATa/MATα ura3/ura3, lys2/lys2, ho::LYS2/ho::LYS2 leu2-K/leu2-R, arg4-nsp,bgl/arg4-nsp,bgl, SynIV-3C-URA3/native* | SK1 | This study |
| ORT4601 | *MATα, ura3, lys2, ho::LYS2, leu2-K, arg4-nsp,bgl* | SK1 | (Sollier *et al*, 2004) |

### Design principles of Syn-3C chromosomes

We aimed at modifying the native sequence of a budding yeast chromosome according to our design principles while introducing as little modifications as possible. Because we were planning on re-assembling only a 150kb window within the genome, we scanned through the overall sequence using a scoring quality function to look for the candidate regions qualifying as the ideal target, i.e. where our principles would introduce a minimal number of mutations. The starting material was the *S. cerevisiae* SK1 strain genome sequence and annotations (Liti *et al*, 2009) and a list of 9 restriction enzymes (EcoRI, HindIII, NdeI, PstI, SacI, SacII, SalI, XbaI, XhoI and DpnII). RE were selected based on their low cost and restriction efficiency. A genome index file was then computed, that contained the following information for each base pair:

- Whether it consists of a “forbidden mutation” site, defined by us as follow: i) start and stop codons of known ORFs, ii) regulatory transcription pre-initiation complexes binding regions identified through ChIP-Seq exo, encompassing TATA-box binding sites (Rhee & Pugh, 2012), iii) the consensus sequence of Autonomous Replicating Sequences (ARS), i.e. the core sequence within *S. cerevisiae* replication origins (list of ARS obtained from oridb (Siow *et al*, 2012), iv) intron borders, v) centromeres, vi) tRNA.
- Whether the position belongs to a restriction site.
- If it belongs to an intergenic or coding region, and in the latter case, the codon it belongs to and its position.

Sliding windows of 150 kb moving with 10kb steps were then generated over the entire genome. In parallel, we defined the restriction pattern we wanted to generate:

- Regularly spaced intervals for 400, 1,500, 2,000 and 6,000 bp
- Gene promoter/terminator (substitutions within a coding sequence strongly preferred) For each window, we computed all possible changes to apply to the genome so that all combinations of five out of the eight chosen 6-cutter enzymes were repositioned to generate all expected new restriction patterns. For each combination of 5 enzymes, all sites were first removed from the genome before being reintroduced at ideal positions. A margin of error in the positioning of the “ideal” position was tolerated (10% of the window size) to maximize the probability of introducing only synonymous mutations within the coding sequence. Once a RS was positioned, the position of the adjacent RS was adjusted based on the newly positioned site so that overall, the distribution of RFs remains as close as possible to the theoretical distribution. Overall, for each enzyme, a quality score was computed for each window based on the difference between the expected distribution of the site, and the real distribution. For each combination of enzyme, a global score corresponding to the sum of the individual scores of each enzyme was computed (see **Fig S1** for schema).

Overall, we selected the 10 “best” windows located at least at 150 kb from either a centromere or a telomere. The quality score was weighted by the presence of “forbidden positions” within the window, for instance when a start codon overlaps a restriction site to be deleted. Finally, a manual curation, aiming at fixing potential conflicts (such as 2 RSs overlapping the same bases, or accidental re-creation of a RS of one enzyme when processing a second one), followed, and was performed on the genome windows presenting the best quality scores.

We chose the final window based also on our research interests, i.e. containing at least two early replicating replication origins (Siow *et al*, 2012; Raghuraman *et al*, 2001), and several hotspots of meiotic DNA double-strand breaks (Pan *et al*, 2011). We also attempted to avoid too many retrotransposable elements or other DNA repeats. The final window was positioned on chromosome IV::700,000-850,000, with restriction patterns as follow: DnpII ↔ 400 bp window; Xbal ↔ 1,500 bp window; HindIII ↔ 2,000 bp window; NdeI ↔ 6,000 bp window; Hhal ↔ promoter/terminator (see summary on **Fig EV2**). 1037 mutations were present in the sequence, the vast majority corresponding to the modifications necessary to reorganize DpnII RS (**Table 1**). Overall, 1037 mutations were introduced, corresponding to 0.7% divergence.

Other modifications were introduced into the sequence. First, PCRTags similar to those used in the Sc2.0 design(Annaluru *et al*, 2014; Dymond *et al*, 2011) specific to either the native or synthetic sequence were also introduced within the window. Performing PCR using these primers allow testing for the presence and absence of the synthetic and native sequence, respectively. PCRTags were manually curated to adapt them to the restriction design, and overall 59 PCR tags out of 154 needed to be modified accordingly.

Overall, a total of 3,229 bp were modified (~2% of the 150,000 bp window). 743 codons were modified, but no change in the sequences of the corresponding proteins were introduced. Although we took great care in the design of the sequence and algorithm, our ongoing experiments nevertheless suggest this design is perfectible and could be simplified. First, windows of 400 and 1,000 bp are probably sufficient to assay the structure at a reasonable resolution. Also, SNPs have to be introduced in repeats or low complexity DNA to facilitate mapping of the reads. Finally, a possibility to find the best frame for each interval/pattern is to start with the positions of RS overlapping forbidden sites as seed.

### Assembly of the redesigned chromosome

The redesigned sequence was split into 52 fragments of ~3,000 bp (i.e., blocks), with 200 bp overlaps between them. In addition sequences corresponding either to the auxotrophic marker genes *URA3* or *LEU2* were added to blocks 20, 37, 52 (*URA3)* and blocks 11, 28, 47 (*LEU2)*, followed by 200 bp sequences of the WT neighboring chromosomal region. The replacement of the native sequence of strain BY4742 with the redesigned blocks was performed through a succession of six transformations, up to 11 blocks at a time (Muller *et al*, 2012).

After each transformation, independent colonies were sampled and PCRs performed at the PCR tags positions to identify the transformants that have replaced all of the native sequence with the redesigned one (**Fig. EV3**). Upon the last transformation, the selected transformant genome was sequenced and the region 707,556-852,114 (144,558 bp) was found to be replaced by the synthetic blocks.

### Growth rate analysis

Growth assays were performed to see if the transformants exhibited changes in fitness. Little to no growth defect could be identified when blocks 1 to 47 replaced the native sequence. The final transformation with blocks 48 till 52 led repeatedly to the recovery of transformants exhibiting a slow-growth, petite phenotype (Slonimski, 1949), reflecting a block in the aerobic respiratory chain pathway and a decrease in ATP. Since the region concerned by the 6^th^ transformation only involved a few kb, we decided to keep the first 145 kb already successfully reassembled and discard this last step. However, crossing this petite strain with a WT strain gave diploids without growth defects. Sporulation of these diploids gave offsprings with growth rates also similar to WT, suggesting stable complementation of mitochondrial genomes.

### Pre-growth and sporulation of yeast

Pre-growth and sporulation of the RSG_Y190 strain was carried out as described (Oh *et al*, 2009). Briefly, cells patched on YPG-Agar plates (1% Yeast Extract, 2% Peptone, 1.5% Agar, 2% Glycerol) from -80°C stocks were streaked on YPD plates (1% Yeast Extract, 2% Peptone, 1.5% Agar, 2% D-glucose, 0.004% Adenine). A single colony was used to inoculate 5 mL YPD liquid culture and grown at 30°C up to saturation. The saturated culture was used to inoculate 350 mL of a freshly made (less than 48 hrs) pre-sporulation media (SPS; 0.5% Yeast extract, 1% Peptone, 1% Potassium Acetate, 1% Ammonium Sulfate, 0.5% Potassium Hydrogen Pthalate, 0.17% Yeast Nitrogen Base lacking all amino-acids, two drops of anti-foaming agent) and grown with robust agitation (320 rpm) in 5L baffled flasks at 30°C. Cells were washed, transferred in 500 mL of sporulation media (SPM; 1% Potassium Acetate, 0.2X of Uracil, Arginine and Leucine, two drops of anti-foaming agent) and put back with robust agitation at 30°C.

### RNA Isolation from Yeast for RNA Sequencing

Three independent RNA-seq libraries were generated for BY4742, SK1 and Syn3C strains. For each library, a single colony was grown in a 2 mL YPD culture overnight at 30°C. The next morning, 10 mL cultures in YPD were started from 10^6^ cells/mL until they reached 2.10^7^ cell/mL. The cells were pelleted by spinning at 5000 rpm at 4°C for 5 min. The pellet was resuspended in 0.5 mL of Tris-HCl (10 mM, pH 7,5) and transferred to a microfuge tube. The cells were pelleted again by spinning briefly and discarding the supernatant. The cells were resuspended in 400 μL RNA TES buffer (10 mM Tris-HCl, pH 7.5 10 mM EDTA 0.5% SDS). 400 μL of acid phenol/chloroform was then added to the cells and vortexed for 1 min, and heated at 65°C for 30 minutes, briefly vortexing some time to time. The cells were placed on ice for 5 min band centrifuge at 13000 rpm at 4°C for 5 min. The aqueous layer (~400 μL) was then transferred into another microfuge tube; an equal amount of phenol/chloroform acid was added a second time, mixed well and centrifuged at 13000 rpm at 4°C for 5 min. The RNA (~400 μL) was precipitated by adding 40 μL of sodium acetate (3 M) and 1,1 mL of absolute ethanol and incubating the tube at -80°C for a least 30 min. The RNA was pelleted by centrifuging at 13000 rpm at 4°C for 20 min. The RNA pellet was then washed with 500 μL of 70% ethanol, air-dried and then resuspended in 50 μL of water. 15 μg were treated with 2U of DNase TURBO (Invitrogen) and cleaned up by phenol extraction and ethanol precipitation before being prepared for sequencing.

### RNA-Seq Analysis of synIII

Single-end non-strand-specific RNA-seq of the Syn3C and BY4742 were performed using Illumina Nextseq and standard TruSeq preparations kits, after depletion of ribosomal RNA. Reads were mapped using Bowtie2 to the reference *S. cerevisiae* BY4742 and Syn3C genome. For each gene, reads were counted if mapping quality was lower than 30 and analyzed for differential expression using DESeq2, with standard parameters.

### Hi-C experiments

*S. cerevisiae* G1 daughter cells of the redesigned strain were recovered from an exponentially growing population through an elutriation procedure (Marbouty *et al*, 2014). Hi-C libraries were generated as described (Lazar-Stefanita *et al*, 2017; Dekker *et al*, 2002). G1 daughter cells were cross-linked for 20 minutes with fresh formaldehyde (3% final concentration). To generate the libraries with different restriction enzymes, aliquots of 3 x 10^9^ cells were resuspended in 10 ml sorbitol 1M and incubated 30 minutes with DTT 5mM and Zymolyase 100T (C_Final_=1 mg/ml) to digest the cell wall. Spheroplasts were then washed first with 5 ml of sorbitol 1M, then with 5 ml of 1X restriction buffer (depending on the restriction enzyme used). The spheroplasts were then resuspended either in 3.5 ml of the corresponding restriction buffer (NEB). For each aliquot/experiment, the cells were then split into three tubes (V=500μL) and incubated in SDS (3%) for 20 minutes at 65°C.

Crosslinked DNA was digested at 37°C overnight with 15 units of the appropriate restriction enzyme (NEB, DpnII, HindIII or NdeI). The digestion mix was then centrifuged for 20 minutes at 18000 g and the supernatant discarded. The pellets were then resuspended and pooled into 400 μL of cold water. Depending on the sequence of the restriction site overhangs, the extremities of the fragments were repaired in the presence of either 14-dCTP biotin or 14-dATP biotin (Invitrogen). Biotinylated DNA molecules were then incubated 4 hours at 16°C in presence of 250 U of T4 DNA ligase (Thermo Scientific, 12.5 ml final volume). DNA purification was achieved through an overnight incubation at 65°C in presence of 250μg/ml proteinase K in 6.2mM EDTA followed by precipitation step in presence of RNAse.

The resulting 3C libraries were sheared and processed into Illumina libraries using custom-made versions of the Illumina PE adapters (Paired-End DNA sample Prep Kit – Illumina – PE-930-1001). Fragments of sizes between 400 and 800 bp were purified using a PippinPrep apparatus (SAGE Science), PCR amplified, and paired-end (PE) sequenced on an Illumina platform (HiSeq2000; 2 x 75 bp).

The accession number for the data reported in this paper is [Database]: [xxxx] (under completion).

### Processing of the reads and contact maps generations

The raw data from each 3C experiment was processed as follow: first, PCR duplicates were collapsed using the 6 Ns present on each of the custom-made adapter and trimmed. Reads were then aligned using Bowtie 2 in its most sensitive mode against *S. cerevisiae* reference genome (native genome) or against the *S. cerevisiae* reference adapted for the Syn-3C region on chromosome 4 (SynIV-3C genome). An iterative alignment procedure was used: for each read the length of the sequence mapped increases gradually from 20 bp until the mapping became unambiguous (mapping quality > 30). Paired reads were aligned independently and each mapped read was assigned to a restriction fragment. Religation events has been filtered out through the information about the orientation of the sequences as described in(Cournac *et al*, 2012). The distribution of the reads along the synthetic region and its native counterpart is represented in **Fig 2**.

Contact matrices were built for the wild type and the mutant by binning the aligned reads into units of single restriction fragments. DpnII and HindIII contact maps for the SynIV-3C region and its native counterpart were randomly resampled in order to present the same number of contacts. The raw contact maps were then subsequently binned into units (i.e. bins) of 600, 1,200, 2,400, 4,800 and 9,600 base pairs. Contacts maps were generated using the *levelplot* function of the R *lattice* package. Matrices for the synthetic region were subsequently obtained by extracting the diagonal blocks for bins falling in the 719,756bp to 849,206bp interval. Outliers has been removed from the matrices if the number of the contacts surpassed by 20 times the top 5‰ threshold of the number of contacts between restriction fragment pairs.

### Statistical analysis

The CV is defined as the ratio between the standard deviation and the mean of the contact histograms at fixed distance s; to take into account the finite-size effect, we discarded bins at the edge of the contact matrix in order to keep the statistics (number of bins) for different values of s constant, up to s < 15,000 bp in DpnII and s < 70,000 bp in HindIII datasets. To show that the improvement is specific to the new restriction pattern and is unlikely to be find spontaneously within the genome, we compared the SynIV-3C results with seven regions of similar size along chromosome 4 (460,856-590,306 bp; 590,306-719,756 bp; 849,206-978,656 bp; 978,656-1,108,106 bp;1,108,106-1,237,556 bp;1,2375,56-1,367,006 bp; 1,367,006-1,496,456 bp). The quality improvement was assessed by computing the logarithm of the ratio of the CVs of the SynIV-3C and native region (**Fig EV5**).

### Southern blot analysis of crossing over formation at the CCT6 hotspot

The *CCT6* locus was chosen because it is the strongest Spo11-induced DSB hotspot in the synthetic region (Pan *et al*, 2011). Cells were harvested and DNA was extracted as described (Oh *et al*, 2009), except that no crosslinking step was performed. The DNA was digested with NdeI and XbaI (New England Biolabs), migrated on a 1% UltraPure Agarose (Invitrogen) 1X TAE for 15 hours at 70 V, and capillary transferred onto a Hybond-XL membrane (GE Healthcare) following the manufacturer instructions. Southern blot hybridization was performed at 65°C in Church buffer (1% BSA, 0.25 M Na2HPO4 pH 7.3, 7% SDS, 1 mM EDTA) with a 1,104 bp-long radiolabeled probe corresponding to the rightmost region common to both the native and SynIV-3C restriction fragments (obtained from SK1 genomic DNA with primers 5’- TGGTGAAGAACTCAGGATTC-3’ and 5’-CAGTTACAATGAAGTCCAGG-3’) and radiolabeled phage lambda DNA (molecular ladder). Radiolabeling was performed with P^32^-αdCTP with the High Prime labeling kit (Roche) following the manufacturer instructions. The membrane was washed, exposed O/N, and the storage Phosphor Screen (GE Healthcare) was scanned on a Typhoon phosphorimager (Molecular Dynamics). The length of the native and SynIV-3C parental fragments are 5,135 bp and 6,157 bp, respectively. CO formation generates two recombinants of 4,453 bp and 6,839 bp. Quantifications were performed under ImageQuant 5.2 (Molecular Dynamics).

## Acknowledgements

We thank Jef Boeke for the Sc2.0 PCRTags sequences for chromosome 4 and for fruitful discussions and comments on the manuscript. We thank Elodie Pirayre and Ivan Moszer for contributing to the initial steps of the design of the algorithm, and Nancy Kleckner, Gianni Liti, Axel Cournac, Martial Marbouty, Luciana Lazar-Stefanita for discussions. Vittore Scolari and Héloïse Muller are recipients of Pasteur-Roux-Cantarini fellowships. This research was supported by funding to R.K. from the European Research Council under the 7th Framework Program (FP7/2007-2013, ERC grant agreement 260822), from ERASynBio and Agence Nationale pour la Recherche (IESY ANR-14-SYNB-0001-03), and to R.K. O.E. and B.L. from Agence Nationale pour la Recherche (MeioRec ANR-13-BSV6-0012).

## Author contributions

HM, SDD, BL, OE, NA, GF and RK designed the sequence. HM assembled the chromosome and performed the Hi-C experiments with GM. AP analyzed meiotic recombination. VS and JM analyzed the data, with contribution from LLS. RK conceived the study and wrote the manuscript, with contributions from JM, VS, HM, AP, and BL.

## References

Acquaviva L, Székvölgyi L, Dichtl B, Dichtl BS, de La Roche Saint André C, Nicolas A & Géli V (2013) The COMPASS subunit Spp1 links histone methylation to initiation of meiotic recombination. Science 339: 215–218

Annaluru N, Muller H, Mitchell LA, Ramalingam S, Stracquadanio G, Richardson SM, Dymond JS, Kuang Z, Scheifele LZ, Cooper EM, Cai Y, Zeller K, Agmon N, Han JS, Hadjithomas M, Tullman J, Caravelli K, Cirelli K, Guo Z, London V, et al (2014) Total Synthesis of a Functional Designer Eukaryotic Chromosome. Science 344: 55–58

Benders GA, Noskov VN, Denisova EA, Lartigue C, Gibson DG, Assad-Garcia N, Chuang R-Y, Carrera W, Moodie M, Algire MA, Phan Q, Alperovich N, Vashee S, Merryman C, Venter JC, Smith HO, Glass JI & Hutchison CA (2010) Cloning whole bacterial genomes in yeast. Nucleic Acids Res. 38: 2558–2569

Blat Y, Protacio RU, Hunter N & Kleckner N (2002) Physical and Functional Interactions among Basic Chromosome Organizational Features Govern Early Steps of Meiotic Chiasma Formation. Cell 111: 791–802

Cournac A, Marie-Nelly H, Marbouty M, Koszul R & Mozziconacci J (2012) Normalization of a chromosomal contact map. BMC Genomics 13: 436

Dekker J, Rippe K, Dekker M & Kleckner N (2002) Capturing chromosome conformation. Science 295: 1306–1311

Dixon JR, Selvaraj S, Yue F, Kim A, Li Y, Shen Y, Hu M, Liu JS & Ren B (2012) Topological domains in mammalian genomes identified by analysis of chromatin interactions. Nature 485: 376–380

Dymond JS, Richardson SM, Coombes CE, Babatz T, Müller H, Annaluru N, Blake WJ, Schwerzmann JW, Dai J, Lindstrom DL, Boeke AC, Gottschling DE, Chandrasegaran S, Bader JS & Boeke JD (2011) Synthetic chromosome arms function in yeast and generate phenotypic diversity by design. Nature 477: 471–476

Glynn EF, Megee PC, Yu H-G, Mistrot C, Unal E, Koshland DE, DeRisi JL & Gerton JL (2004) Genome-Wide Mapping of the Cohesin Complex in the Yeast Saccharomyces cerevisiae. PLOS Biol. 2: e259

Hughes JR, Roberts N, McGowan S, Hay D, Giannoulatou E, Lynch M, De Gobbi M, Taylor S, Gibbons R & Higgs DR (2014) Analysis of hundreds of cis-regulatory landscapes at high resolution in a single, high-throughput experiment. Nat. Genet. 46: 205–212

Hunter N & Kleckner N (2001) The Single-End Invasion: An Asymmetric Intermediate at the Double-Strand Break to Double-Holliday Junction Transition of Meiotic Recombination. Cell 106: 59–70

Imakaev M, Fudenberg G, McCord RP, Naumova N, Goloborodko A, Lajoie BR, Dekker J & Mirny LA (2012) Iterative correction of Hi-C data reveals hallmarks of chromosome organization. Nat. Methods 9: 999–1003

Ito M, Kugou K, Fawcett JA, Mura S, Ikeda S, Innan H & Ohta K (2014) Meiotic recombination cold spots in chromosomal cohesion sites. Genes Cells Devoted Mol. Cell. Mech. 19: 359–373

Kim KP, Weiner BM, Zhang L, Jordan A, Dekker J & Kleckner N (2010) Sister cohesion and structural axis components mediate homolog bias of meiotic recombination. Cell 143: 924–937

Kleckner N, Zickler D, Jones GH, Dekker J, Padmore R, Henle J & Hutchinson J (2004) A mechanical basis for chromosome function. Proc. Natl. Acad. Sci. U. S. A. 101: 12592–12597

Klein F, Mahr P, Galova M, Buonomo SBC, Michaelis C, Nairz K & Nasmyth K (1999) A Central Role for Cohesins in Sister Chromatid Cohesion, Formation of Axial Elements, and Recombination during Yeast Meiosis. Cell 98: 91–103

Koszul R (2016) Beyond the bounds of evolution: Synthetic chromosomes… How and what for? C. R. Biol. Available at: http://dx.doi.org/10.1016/j.crvi.2016.05.007

Koszul R, Kim KP, Prentiss M, Kleckner N & Kameoka S (2008) Meiotic Chromosomes Move by Linkage to Dynamic Actin Cables with Transduction of Force through the Nuclear Envelope. Cell 133: 1188–1201

Koszul R & Kleckner N (2009) Dynamic chromosome movements during meiosis: a way to eliminate unwanted connections? Trends Cell Biol. 19: 716–724

Lajoie BR, Dekker J & Kaplan N (2015) The Hitchhiker’s guide to Hi-C analysis: practical guidelines. Methods San Diego Calif 72: 65–75

Lazar-Stefanita L, Scolari VF, Mercy G, Muller H, Guérin TM, Thierry A, Mozziconacci J & Koszul R (2017) Cohesins and condensins orchestrate the 4D dynamics of yeast chromosomes during the cell cycle. EMBO J.: e201797342

Le TBK, Imakaev MV, Mirny LA & Laub MT (2013) High-resolution mapping of the spatial organization of a bacterial chromosome. Science 342: 731–734

Lesne A, Riposo J, Roger P, Cournac A & Mozziconacci J (2014) 3D genome reconstruction from chromosomal contacts. Nat. Methods 11: 1141–1143

Lieberman-Aiden E, Berkum NL van, Williams L, Imakaev M, Ragoczy T, Telling A, Amit I, Lajoie BR, Sabo PJ, Dorschner MO, Sandstrom R, Bernstein B, Bender MA, Groudine M, Gnirke A, Stamatoyannopoulos J, Mirny LA, Lander ES & Dekker J (2009) Comprehensive Mapping of Long-Range Interactions Reveals Folding Principles of the Human Genome. Science 326: 289–293

Liti G, Carter DM, Moses AM, Warringer J, Parts L, James SA, Davey RP, Roberts IN, Burt A, Koufopanou V, Tsai IJ, Bergman CM, Bensasson D, O’Kelly MJT, van Oudenaarden A, Barton DBH, Bailes E, Nguyen AN, Jones M, Quail MA, et al (2009) Population genomics of domestic and wild yeasts. Nature 458: 337–341

Marbouty M, Ermont C, Dujon B, Richard G-F & Koszul R (2014) Purification of G1 daughter cells from different Saccharomycetes species through an optimized centrifugal elutriation procedure. Yeast 31: 159–166

Marie-Nelly H, Marbouty M, Cournac A, Liti G, Fischer G, Zimmer C & Koszul R (2014) Filling annotation gaps in yeast genomes using genome-wide contact maps. Bioinforma. Oxf. Engl. 30: 2105–2113

Mercy G, Mozziconacci J, Scolari VF, Yang K, Zhao G, Thierry A, Luo Y, Mitchell LA, Shen M, Shen Y, Walker R, Zhang W, Wu Y, Xie Z, Luo Z, Cai Y, Dai J, Yang H, Yuan Y-J, Boeke JD, et al (2017) 3D organization of synthetic and scrambled chromosomes. Science 355: eaaf4597

Muller H, Annaluru N, Schwerzmann JW, Richardson SM, Dymond JS, Cooper EM, Bader JS, Boeke JD & Chandrasegaran S (2012) Assembling large DNA segments in yeast. Methods Mol. Biol. Clifton NJ 852: 133–150

Naumova N, Imakaev M, Fudenberg G, Zhan Y, Lajoie BR, Mirny LA & Dekker J (2013) Organization of the mitotic chromosome. Science 342: 948–953

Nora EP, Lajoie BR, Schulz EG, Giorgetti L, Okamoto I, Servant N, Piolot T, van Berkum NL, Meisig J, Sedat J, Gribnau J, Barillot E, Blüthgen N, Dekker J & Heard E (2012) Spatial partitioning of the regulatory landscape of the X-inactivation centre. Nature 485: 381–385

Oh SD, Jessop L, Lao JP, Allers T, Lichten M & Hunter N (2009) Stabilization and Electrophoretic Analysis of Meiotic Recombination Intermediates in *Saccharomyces cerevisiae*. In Meiosis pp 209–234. Humana Press Available at: https://link.springer.com/protocol/10.1007/978-1-59745-527-5_14 [Accessed December 14, 2017]

Padmore R, Cao L & Kleckner N (1991) Temporal comparison of recombination and synaptonemal complex formation during meiosis in S. cerevisiae. Cell 66: 1239–1256

Pan J, Sasaki M, Kniewel R, Murakami H, Blitzblau HG, Tischfield SE, Zhu X, Neale MJ, Jasin M, Socci ND, Hochwagen A & Keeney S (2011) A hierarchical combination of factors shapes the genome-wide topography of yeast meiotic recombination initiation. Cell 144: 719–731

Raghuraman MK, Winzeler EA, Collingwood D, Hunt S, Wodicka L, Conway A, Lockhart DJ, Davis RW, Brewer BJ & Fangman WL (2001) Replication dynamics of the yeast genome. Science 294: 115–121

Rao SSP, Huntley MH, Durand NC, Stamenova EK, Bochkov ID, Robinson JT, Sanborn AL, Machol I, Omer AD, Lander ES & Aiden EL (2014) A 3D map of the human genome at kilobase resolution reveals principles of chromatin looping. Cell 159: 1665–1680

Rhee HS & Pugh BF (2012) Genome-wide structure and organization of eukaryotic pre-initiation complexes. Nature 483: 295–301

Sexton T, Yaffe E, Kenigsberg E, Bantignies F, Leblanc B, Hoichman M, Parrinello H, Tanay A & Cavalli G (2012) Three-dimensional folding and functional organization principles of the Drosophila genome. Cell 148: 458–472

Siow CC, Nieduszynska SR, Müller CA & Nieduszynski CA (2012) OriDB, the DNA replication origin database updated and extended. Nucleic Acids Res. 40: D682–D686

Sollier J, Lin W, Soustelle C, Suhre K, Nicolas A, Géli V & Saint-André C de LR (2004) Set1 is required for meiotic S-phase onset, double-strand break formation and middle gene expression. EMBO J. 23: 1957–1967

Sommermeyer V, Béneut C, Chaplais E, Serrentino ME & Borde V (2013) Spp1, a Member of the Set1 Complex, Promotes Meiotic DSB Formation in Promoters by Tethering Histone H3K4 Methylation Sites to Chromosome Axes. Mol. Cell 49: 43–54

Trelles-Sticken E, Loidl J & Scherthan H (1999) Bouquet formation in budding yeast: initiation of recombination is not required for meiotic telomere clustering. J. Cell Sci. 112: 651–658

Zickler D & Kleckner N (1999) Meiotic chromosomes: integrating structure and function. Annu. Rev. Genet. 33: 603–754

Zickler D & Kleckner N (2016) A Few of Our Favorite Things: Pairing, the Bouquet, Crossover Interference and Evolution of Meiosis. Semin. Cell Dev. Biol. 54: 135–148

